# Co-linear Chaining with Overlaps and Gap Costs

**DOI:** 10.1101/2021.02.03.429492

**Authors:** Chirag Jain, Daniel Gibney, Sharma V. Thankachan

## Abstract

Co-linear chaining has proven to be a powerful heuristic for finding near-optimal alignments of long DNA sequences (e.g., long reads or a genome assembly) to a reference. It is used as an intermediate step in several alignment tools that employ a seed-chain-extend strategy. Despite this popularity, efficient subquadratic-time algorithms for the general case where chains support anchor overlaps and gap costs are not currently known. We present algorithms to solve the co-linear chaining problem with anchor overlaps and gap costs in *Õ*(*n*) time, where *n* denotes the count of anchors. We also establish the first theoretical connection between co-linear chaining cost and edit distance. Specifically, we prove that for a fixed set of anchors under a carefully designed chaining cost function, the optimal ‘anchored’ edit distance equals the optimal co-linear chaining cost. Finally, we demonstrate experimentally that optimal co-linear chaining cost under the proposed cost function can be computed orders of magnitude faster than edit distance, and achieves correlation coefficient above 0.9 with edit distance for closely as well as distantly related sequences.

## 1 Introduction

Computing an optimal alignment between two sequences is one of the most fundamental problems in computational biology. Unfortunately, conditional lower-bounds suggest that an algorithm for computing an optimal alignment, or edit distance, in strongly subquadratic time is unlikely [3,10]. This lower-bound indicates a challenge for scaling the computation of edit distance to high-throughput sequencing data. Instead, heuristics are often used to obtain an approximate solution in less time and space. One such popular heuristic is co-linear chaining. This technique involves precomputing fragments between the two sequences that closely agree (in this work, exact matches called *anchors*), then determining which of these anchors should be kept within the alignment (see Fig. 1). Techniques along these lines are used in long-read mappers [6,11,14,15,23,24,26] and generic sequence aligners [2,5,13,18,22]. We will focus on the following problem (described formally in Section 2): Given a set of *n* anchors, determine an optimal ordered subset (or chain) of these anchors.

**Fig. 1:**
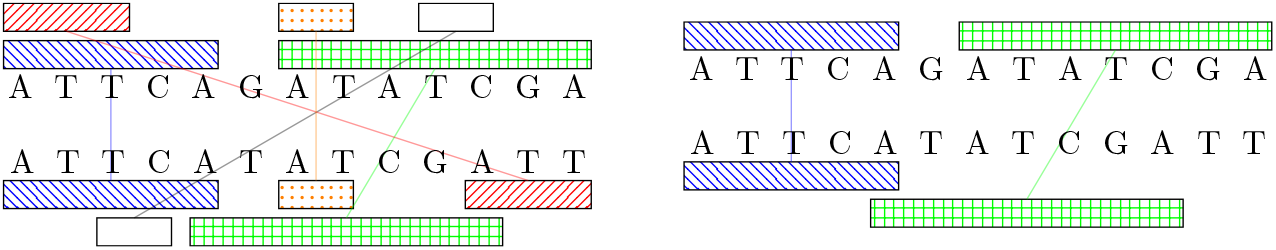
(Left) Anchors representing a set of exact matches are shown as rectangles. The co-linear chaining problem is to find an optimal ordered subset of anchors subject to some cost function. (Right) A chain of overlapping anchors.

Several algorithms have been developed for the co-linear chaining [1,16,27,30] and even more in the context of sparse dynamic programming [8,9,19,21,32,17]. Solutions with different time complexities exist for different variations of this problem. These depend on the cost-function assigned to a chain and the types of chains permitted. Solutions include an algorithm running in *O*(*n* log *n* log log *n*) time for a simpler variant of the problem where anchors used in a solution must be non-overlapping [1]. More recently, Mäkinen and Sahlin gave an algorithm running in *O*(*n* log *n*) time where anchor overlaps are allowed, but gaps between anchors are not considered in the cost-function [16]. None of the solutions introduced thus far provide a subquadratic time algorithm for variations that use both overlap and gap costs. However, including overlaps and gaps into a cost-function is a more realistic model for anchor chaining. For example, consider a simple scenario where minimizers [25] are used to identify anchors. Suppose query and reference sequences are identical, then adjacent minimizer-anchors will likely overlap. Not allowing anchor overlaps during chaining will lead to a penalty cost associated with gaps between chained anchors despite the two strings being identical. Therefore, depending on the type of anchor, there may be no reason to assume that in an optimal alignment the anchors would be non-overlapping. At the same time, not penalizing long gaps between the anchors is unlikely to produce correct alignments. This is why both anchor overlaps and gap costs are supported during chaining in widely-used aligners, e.g., Minimap2 [12,14], Nucmer4 [18]. This work’s contribution is the following:

– We provide the first algorithm running in subquadratic, *Õ*(*n*) time for chaining with overlap and gap costs^1^. Refinements based on the specific type of anchor and chain under consideration are also given. These refinements include an *O*(*n* log^2^ *n*) time algorithm for the case where all anchors are of the same length, as is the case with *k*-mers.
– When *n* is not too large (less than the sequence lengths), we present an algorithm with *O*(*n* · *OPT* + *n* log *n*) average-case time where *OPT* is the optimal solution value. This provides a simple algorithm that is efficient in practice.
– Using a carefully designed cost-function, we mathematically relate the optimal chaining cost with a generalized version of edit distance, which we call *anchored edit distance*. This is equivalent to the usual edit distance with the modification that matches performed without the support of an anchor have unit cost. A more formal definition appears in Section 2. With our cost function, we prove that the optimal chaining cost is equal to the anchored edit distance.
– We empirically demonstrate that computing optimal chaining cost is orders of magnitude faster than computing edit distance, especially in semi-global comparison mode. We also demonstrate a strong correlation between optimal chaining cost and edit distance. The correlation coefficients are favorable when compared to suboptimal chaining methods implemented in Minimap2 and Nucmer4.

## 2 Concepts and Definitions

Let *S*_1_ and *S*_2_ be two strings of lengths |*S*_1_| and |*S*_2_| respectively. An anchor interval pair ([*a*..*b*], [*c*..*d*]) signifies an exact match between *S*_1_[*a*..*b*] and *S*_2_[*c*..*d*]. For an anchor *I*, we denote these values as *I*.*a, I*.*b, I*.*c*, and *I*.*d*. Here *b* − *a* = *d* − *c* and *S*_1_[*a* + *j*] = *S*_2_[*c* + *j*] for all 0≤ *j*≤ *b* − *a*. We say that the character match *S*_1_[*a* + *j*] = *S*_2_[*c* + *j*], 0 ≤ *j* ≤ *b* − *a*, is *supported* by the anchor ([*a*..*b*], [*c*..*d*]). Maximal exact matches (MEMs), maximal unique matches (MUMs), or *k*-mer matches are some of the common ways to define anchors. Maximal unique matches [7] are a subset of maximal exact matches, having the added constraint that the pattern involved occurs only once in both strings. If all intervals across all anchors have the same length (e.g., using *k*-mers), we say that the *fixed-length* property holds.

Our algorithms will make use of dynamic range minimum queries (RmQs). For a set of *n d*-dimensional points, each with an associated weight, a ‘query’ consists of an orthogonal *d*-dimensional range. The query response is the point in that range with the smallest weight. Using known techniques in computational geometry, a data structure can be built in *O*(*n* log^*d*−1^ *n*) time and space, that can both answer queries and modify a point’s weight in *O*(log^*d*^ *n*) time [4].

### 2.1 Co-linear Chaining Problem with Overlap and Gap Costs

Given a set of *n* anchors 𝒜 for strings *S*_1_ and *S*_2_, we assume that 𝒜 already contains two *end-point* anchors 𝒜_*left*_ = ([0, 0], [0, 0]) and 𝒜_*right*_ = ([|*S*_1_ |+ 1, |*S*_1_|+ 1], [|*S*_2_ |+1, |*S*_2_ |+1]). We define the strict precedence relationship ≺ between two anchors *I*′ := 𝒜 [*j*] and *I* := 𝒜 [*i*] as *I*′ ≺ *I* if and only if *I*′.*a* ≤*I*.*a, I*′.*b* ≤*I*.*b, I*′.*c* ≤*I*.*c, I*′.*d* ≤*I*.*d*, and strict inequality holds for at least one of the four inequalities. In other words, the interval belonging to *I*′ for *S*_1_ (resp. *S*_2_) should start before or at the starting position of the interval belonging to *I* for *S*_1_ (resp. *S*_2_) and should not extend past it. We also define the weak precedence relation ≺_*w*_ as *I*′ ≺_*w*_ *I* if and only if *I*′.*a* ≤*I*.*a, I*′.*c* ≤*I*.*c* and strict inequality holds for at least one of the two inequalities, i.e., intervals belonging to *I*′ should start before or at the starting position of intervals belonging to *I*, but now intervals belonging to *I*′ can be extended past the intervals belonging to *I*. The aim of the problem is to find a totally ordered subset (a chain) of 𝒜 that achieves the minimum cost under the cost function presented next. We specify whether we mean a chain under strict precedence or under weak precedence when necessary.

#### Cost function

For *I*′ ≺*I*, the function *connect*(*I*′, *I*) is designed to indicate the cost of connecting anchor *I*′ to anchor *I* in a chain. The chaining problem asks for a chain of *m* ≤ *n* anchors, 𝒜′[1], 𝒜′[2], …, 𝒜′[*m*], such that the following properties hold: (i) 𝒜′[1] = 𝒜_*left*_, (ii) 𝒜′[*m*] = 𝒜_*right*_, (iii) 𝒜′[1] ≺ *𝒜*′[2] ≺ …≺ 𝒜^′^ [*m*], and (iv) the cost 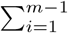 *connect*(𝒜′ [*i*], *𝒜* ′ [*i* + 1]) is minimized.

We next define the function *connect*. In Section 4, we will see that this definition is well motivated by the relationship with anchored edit distance. For a pair of anchors *I*′, *I* such that *I*′ ≺ *I*:

– The gap in string *S*_1_ between anchors *I*′ and *I* is *g*_1_ = max(0, *I*.*a − I*′.*b*− 1). Similarly, the gap between the anchors in string *S*_2_ is *g*_2_ = max(0, *I*.*c* − *I*′.*d* −1). Define the gap cost *g*(*I*′, *I*) = max(*g*_1_, *g*_2_).
– The overlap *o*_1_ is defined such that *I*′.*b* − *o*_1_ reflects the non-overlapping prefix of anchor *I*′ in string *S*_1_. Specifically, *o*_1_ = max(0, *I*′.*b* − *I*.*a* + 1). Similarly, define *o*_2_ = max(0, *I*′.*d* − *I*.*c* + 1). We define the overlap cost as *o*(*I*′, *I*) = |*o*_1_ − *o*_2_ |.
– Lastly, define *connect*(*I*′, *I*) = *g*(*I*′, *I*) + *o*(*I*′, *I*).

The same definitions are used for weak precedence, only using ≺_*w*_ in the place of ≺. Regardless of the definition of *connect*, the above problem can be trivially solved in *O*(*n*^2^) time and *O*(*n*) space. First sort the anchors by the component 𝒜[·].*a* and let 𝒜′ be the sorted array. The chaining problem then has a direct dynamic programming solution by filling an *n*-sized array *C* from left-to-right, such that *C*[*i*] reflects the cost of an optimal chain that ends at anchor 𝒜′[*i*]. The value *C*[*i*] is computed using the recursion: *C*[*i*] = min_𝒜*′*[*k*]≺ 𝒜*′*[*i*]_(*C*[*k*] +*connect* (𝒜′[*k*], *𝒜* ′[*i*]) where the base case associated with anchor 𝒜_*left*_ is *C*[1] = 0. The optimal chaining cost will be stored in *C*[*n*] after spending *O*(*n*^2^) time. We will provide an *O*(*n* log^4^ *n*) time algorithm for this problem.

### 2.2 Anchored Edit Distance

The edit distance problem is to identify the minimum number of operations (substitutions, insertions, or deletions) that must be applied to string *S*_2_ to transform it to *S*_1_. Edit operations can be equivalently represented as an alignment (a.k.a. edit transcript) that specifies the associated matches, mismatches and gaps while placing one string on top of another. The *anchored edit distance problem* is as follows: given strings *S*_1_ and *S*_2_ and a set of *n* anchors 𝒜, compute the optimal edit distance subject to the condition that a match supported by an anchor has edit cost 0, and a match that is not supported by an anchor has edit cost 1.

The above problem is solvable in *O* (|*S*_1_| |*S*_2_|) time and space. We can assume that input does not contain redundant anchors, therefore, the count of anchors is ≤|*S*_1_| |*S*_2_|. Next, the standard dynamic programming recursion for solving the edit distance problem can be revised. Let *D*[*i, j*] denote anchored edit distance between *S*_1_[1, *i*] and *S*_2_[1, *j*], then *D*[*i, j*] = min(*D*[*i* − 1, *j* − 1] + *x, D*[*i* − 1, *j*] + 1, *D*[*i, j* −1] + 1) where *x* = 0 if *S*_1_[*i*] = *S*_2_[*j*] and the match is supported by some anchor, and *x* = 1 otherwise.

### 2.3 Graph Representation of Alignment

It is useful to consider the following representation of an alignment of two strings *S*_1_ and *S*_2_. As illustrated in Figure 2, we have a set of |*S*_1_| top vertices and |*S*_2_| bottom vertices. There are two types of edges between the top and bottom vertices: (i) A solid edge from *i*th top vertex to the *j*th bottom vertex. This represents an anchor supported character match between the *i*th character in *S*_1_ and the *j*th character in *S*_2_; (ii) A dashed edge from the *i*th top vertex to the *j*th bottom vertex. This represents a character being substituted to form a match between *S*_1_[*i*] and *S*_2_[*j*] or a character match not supported by an anchor. All unmatched vertices are labeled with an ‘x’ to indicate that the corresponding character is deleted. An important observation is that no two edges cross.

**Fig. 2:**
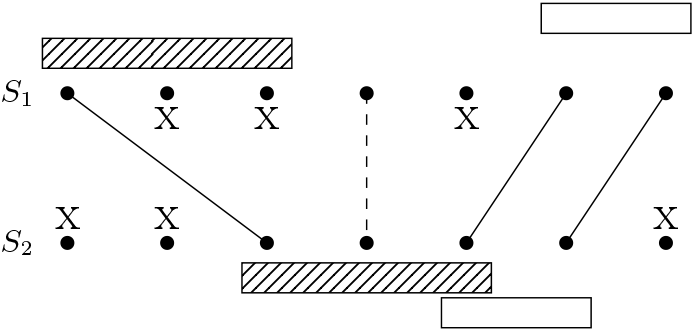
The graph representation of an alignment. Solid edges represent anchor-supported character matches, dashed edges represent substitutions and unsupported matches, and x’s represent deletions. We use ℳ to denote an alignment. Here *EDIT* (ℳ) = 7, the total number of x’s and dashed edges.

In a solution to the anchored edit distance problem every solid edge must be ‘supported’ by an anchor. By ‘supported’ here we mean that the match between the corresponding characters in *S*_1_ and *S*_2_ is supported by an anchor. In Figure 2, these anchors are represented with rectangles above and below the vertices. We use ℳ to denote a particular alignment. We also associate an edit cost with the alignment, denoted as *EDIT* (ℳ). This is equal to the number of vertices marked with x in ℳ plus the number of dashed edges in ℳ.

## 3 Our Algorithms

### Theorem 1.

*The co-linear chaining problem with overlap and gap costs can be solved in time Õ*(*n*). *In particular, in time O*(*n* log^2^ *n*) *for chains with fixed-length anchors; in time O*(*n* log^3^ *n*) *for chains under weak precedence; and in time O*(*n* log^4^ *n*) *for chains under strict precedence*.

The proposed algorithm still uses the recursive formula given in Section 2.1. However, it uses range minimum query (RmQ) data structures to avoid having to check every anchor 𝒜 [*k*] where 𝒜 [*k*].*a* < *𝒜* [*i*].*a*. We achieve this by considering six cases concerning the optimal choice of the prior anchor. We use the best of the six distinct possibilities to determine the optimal *C*[*i*] value. This *C*[*i*] value is then used to update the RmQ data structures. For the strict precedence case, some of the six cases require up to four dimensions for the range minimum queries. When only weak precedence is required, we reduce this to at most three dimensions. When the fixed-length property holds (e.g., *k*-mers), we reduce this to two dimensions.

### Algorithm for chains under strict precedence

The first step is to sort the set of anchors 𝒜 using the key 𝒜[·].*a*. Let 𝒜′ be the sorted array. We will next use six RmQ data structures labeled 𝒯_1*a*_, 𝒯_1*b*_, 𝒯_2*a*_, 𝒯_2*b*_, 𝒯_3*a*_, 𝒯_3*b*_. These RmQ data structures are initialized with the following points for every anchor: For anchor *I* ∈ *𝒜* ′: 𝒯_1*a*_ is initialized with the point (*I*.*b, I*.*d* − *I*.*b*), 𝒯_1*b*_ with (*I*.*d, I*.*d* − *I*.*b*), 𝒯_2*a*_ with (*I*.*b, I*.*c, I*.*d*), 𝒯_2*b*_ with (*I*.*b, I*.*d*), 𝒯_3*a*_ with (*I*.*b, I*.*c, I*.*d, I*.*d* − *I*.*b*), and 𝒯_3*b*_ with (*I*.*b, I*.*d, I*.*d*− *I*.*b*). All weights are initially set to ∞ except for *I* =𝒜 _*left*_, where the corresponding points are given weight 0. We then process the anchors in sorted order and update the RmQ data structures after each iteration. On the *i*th iteration, for *j* < *i*, we let *C*[*j*] be the optimal co-linear chaining cost of any ordered subset of 𝒜′[1], 𝒜′[2], …, 𝒜′[*j*] that ends with 𝒜′[*j*]. For *i* > 1, RmQ queries are used to find the optimal *j* < *i* by considering six different cases. We let *I* = 𝒜′[*i*], *I*′ = 𝒜′[*j*], and *C*[*I*′] = *C*[*j*].

The query for each RmQ structure is determined by the different inequalities relating *I*.*a, I*.*b, I*.*c*, and *I*.*d* to previous anchors in the case considered. For example, in Case 1.a (Figure 3), it can be seen that *I*′.*b* < *I*.*a* and *I*.*a − I*′.*b* < *I*.*c − I*′.*d*, making *I*′.*b* ∈ [0, *I*.*a* −1] and *I*′.*d* − *I*′.*b* ∈ [−∞, *I*.*c* − *I*.*a*], motivating the query input [0, *I*.*a* −1] *×* [−∞, *I*.*c* − *I*.*a*]. At the same time, the values stored in these RmQ structures are determined by the expression for the co-linear chaining cost in that case, *C*[*I*′] + *I*.*c* − *I*′.*d* −1. Note that the values stored in each RmQ structure depend only on previously processed anchors and are combined with the values *I*.*a, I*.*b, I*.*c*, and *I*.*d* for the current anchor *I* being processed to obtain the appropriate cost. Hence, for 𝒯_1*a*_ we store values of the form *C*[*I*′] − *I*′.*d* and combine this with *I*.*c* to obtain the cost. The other cases can be similarly analyzed.

1. Case: *I*′ disjoint from *I*.
  a. Case: The gap in *S*_1_ is less or equal to gap in *S*_2_ (Fig. 3 (Left)). The range minimum query (query input) is of the form: [0, *I*.*a* − 1] *×* [−∞, *I*.*c* − *I*.*a*]. Let the query response (weight) from *T*_1*a*_ be *v*_1*a*_ = min*{C*[*I*′] − *I*′.*d* : (*I*′.*b, I*′.*d*− *I*′.*b*) ∈ [0, *I*.*a*−1]*×*[−∞, *I*.*c*− *I*.*a*]*}* and let *C*_1*a*_ = *v*_1*a*_ +*I*.*c*−1.
  b. Case: The gap in *S*_2_ is less than gap in *S*_1_. The range minimum query is of the form [0, *I*.*c* − 1] *×* [*I*.*c* − *I*.*a*+1, ∞ ]. Let the query response from 𝒯_1*b*_ be *v*_1*b*_ = min {*C*[*I*′] − *I*′.*b* : (*I*′.*d, I*′.*d* − *I*′.*b*) ∈ [0, *I*.*c* − 1]×[*I*.*c I*.*a*+1, ∞ ] and let *C*_1*b*_ = *v*_1*b*_ + *I*.*a* − 1.
2. Case: *I*′ and *I* overlap in only one dimension.
  a. Case: *I*′ and *I* overlap only in *S*_2_ (Fig 3 (Middle)). The range minimum query is of the form [0, *I*.*a* − 1] *×* [0, *I*.*c*] *×* [*I*.*c, I*.*d*]. Let the query response from 𝒯_2*a*_ be *v*_2*a*_ = min {*C*[*I*′] − *I*′.*b* + *I*′.*d* : (*I*′.*b, I*′.*c, I*′.*d*) ∈ [0, *I*.*a* − 1] *×* [0, *I*.*c*] *×* [*I*.*c, I*.*d*] and let *C*_2*a*_ = *v*_2*a*_ + *I*.*a* − *I*.*c*.
  b. Case: *I*′ and *I* overlap only in *S*_1_. Since the anchors are sorted on 𝒜[·].*a*, this can be done with a two dimensional RmQ structure. The range minimum query is of the form [*I*.*a, I*.*b*] *×* [0, *I*.*c* − 1]. Let the query response from 𝒯 _2*b*_ be *v*_2*b*_ = min {*C*[*I*′] + *I*′.*b* − *I*′.*d* : (*I*′.*b, I*′.*d*) ∈ [*I*.*a, I*.*b*] *×* [0, *I*.*c* − 1]} and let *C*_2*b*_ = *v*_2*b*_ + *I*.*c* − *I*.*a*.
3. Case: *I*′ and *I* overlap in both dimensions.
  a. Case: Greater overlap in *S*_2_ (Fig. 3 (Right)). Here, |*o*_1_ − *o*_2_| = *o*_2_ − *o*_1_ = *I*′.*d* − *I*.*c* − (*I*′.*b* − *I*.*a*). The range minimum query is of the form [*I*.*a, I*.*b*]*×*[0, *I*.*c*]*×*[*I*.*c, I*.*d*]*×*[*I*.*c*− *I*.*a*+1, ∞]. Let the query response from 𝒯_3*a*_ be *v*_3*a*_ = min*{C*[*I*′]− *I*′.*b*+*I*′.*d* : (*I*′.*b, I*′*c, I*′.*d, I*′.*d*− *I*′.*b*) ∈ [*I*.*a, I*.*b*]*×* [0, *I*.*c*] *×* [*I*.*c, I*.*d*] *×* [*I*.*c* − *I*.*a* + 1, ∞]*}* and let *C*_3*a*_ = *v*_3*a*_ + *I*.*a* − *I*.*c*.
  b. Case: Greater or equal overlap in *S*_1_. Here, |*o*_1_ − *o*_2_| = *o*_1_ − *o*_2_ = *I*′.*b* − *I*.*a* − (*I*′.*d* − *I*.*c*). If *o*_1_ ≥ *o*_2_ > 0, *I*′.*b* ∈ [*I*.*a, I*.*b*], and *I*′.*a* ∈ [0, *I*.*a*], then *I*′.*c* ∈ [0, *I*.*c*]. Hence, the range minimum query is of the form [*I*.*a, I*.*b*] *×* [*I*.*c, I*.*d*] *×* [−∞, *I*.*c* − *I*.*a*]. Let the query response from 𝒯_3*b*_ be *v*_3*b*_ = min*{C*[*I*′] + *I*′.*b* − *I*′.*d* : (*I*′.*b, I*′.*d, I*′.*d* − *I*′.*b*) ∈ [*I*.*a, I*.*b*] *×* [*I*.*c, I*.*d*] *×* [−∞, *I*.*c* − *I*.*a*]*}* and let *C*_3*b*_ = *v*_3*b*_ − *I*.*a* + *I*.*c*.

**Fig. 3:**
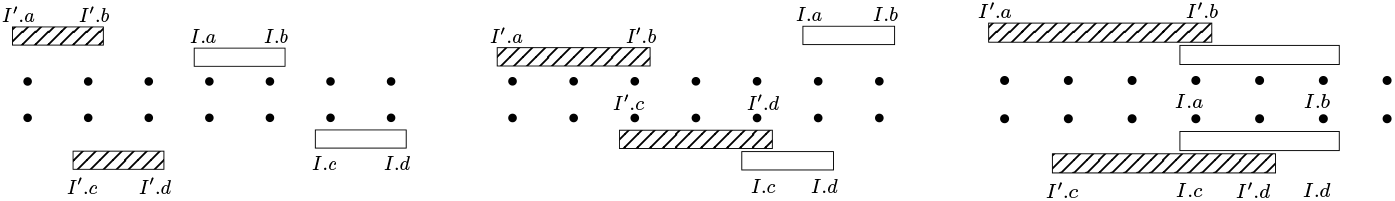
(Left) Case 1.a. Colinear chaining cost is *C*[*I*′] + *g*_2_ = *C*[*I*′] + *I*.*c* − *I*′.*d* − 1. (Middle) Case 2.a. Chaining cost is *C*[*I*′] + *g*_1_ + *o*_2_ = *C*[*I*′] + *I*.*a* − *I*′.*b* + *I*′.*d* − *I*.*c*. (Right) Case 3.a. Chaining cost is *C*[*I*′] + *o*_2_ − *o*_1_ = *C*[*j*] + *I*′.*d* − *I*.*c* − (*I*′.*b* − *I*.*a*).

Finally, let *C*[*i*] = min(*C*_1*a*_, *C*_1*b*_, *C*_2*a*_, *C*_2*b*_, *C*_3*a*_, *C*_3*b*_) and update the RmQ structures as shown in the Pseudo-code in Appendix 1. In the pseudo-code, every RmQ structure 𝒯 has the query method 𝒯.*RmQ*() which takes as arguments an interval for each dimension. It also has the method 𝒯.*update*(), which takes a point and a weight and updates the point to have the new weight. The four-dimensional RmQ structures for Case 3.a require *O*(log^4^ *n*) time per query and update, causing an over time complexity that is *O*(*n* log^4^ *n*). In Appendix 2 we present the modifications for weak precendence and fixed-length anchors.

## 4 Proof of Equivalence

### Theorem 2.

*For a fixed set of anchors 𝒜, the following quantities are equal: the anchored edit distance, the optimal co-linear chaining cost under strict precedence, and the optimal co-linear chaining cost under weak precedence*.

The optimal co-linear chaining cost is defined using the cost function described in Section 2.1. An implication of Theorems 1 and 2 is that if only the anchored edit distance is desired (and not an optimal strictly ordered anchor chain), there exists a *O*(*n* log^3^ *n*) for computing this value.

Theorem 2 will follow from Lemmas 1 and 2.

### Lemma 1.

*Anchored edit distance* ≤ *optimal co-linear chaining cost under weak precedence* ≤ *optimal co-linear chaining cost under strict precedence*.

*Proof*. The second inequality follows from the observation that every set of anchors ordered under strict precedence is also ordered under weak precedence. We now focus on the inequality between anchored edit distance and co-linear chaining cost under weak precedence. Starting with an anchor chain under weak precedence, 𝒜 [1], 𝒜 [2], …with associated co-linear chaining cost *x*, we provide an alignment with an anchored edit distance that is at most *x*. This alignment is obtained using a greedy algorithm that works from left-to-right, always taking the closest exact match when possible, and when not possible, a character substitution or unsupported exact match, or if none of these are possible, a deletion. We now present the details.

### Greedy Algorithm

Assume inductively that all symbols in *S*_1_[1, [*i*].*b*] and *S*_2_[1, *𝒜* [*i*].*d*] have been processed, that is, either matched, substituted, or deleted (represented by check-marks in Figures 4-6). The base case of this induction holds trivially for 𝒜 _*left*_. We consider the anchor 𝒜 [*i* + 1] and the possible cases regarding its position relative to 𝒜 [*i*]. Symmetric cases that only swap the roles of *S*_1_ and *S*_2_ are ignored. To ease notation, let *I*′ = 𝒜 [*i*] and *I* = 𝒜 [*i* + 1].

1. **Case** *I*′.*b* ≥ *I*.*b* and *I*′.*d* ≥ *I*.*c* (Fig 4): To continue the alignment, delete the substring *S*_2_[*I*′.*d*+1, *I*.*d*] from *S*_2_. This has edit cost *I*.*d − I*′.*d*. We can assume both intervals of *I*′ are not nested in intervals of *I*, hence *connect*(*I*′, *I*) = *o*_1_ − *o*_2_ = *I*′.*b* − *I*.*a* − *I*′.*d* + *I*.*c* ≥ *I*.*c* + *I*.*b* − *I*.*a* − *I*′.*d* = *I*.*d* − *I*′.*d*.
2. **Case** *I*′.*b* ≥ *I*.*b* and *I*′.*d* < *I*.*c* (Fig 4): Delete the substring *S*_2_[*I*′.*d* + 1, *I*.*d*] from *S*_2_, with edit cost *I*.*d* − *I*′.*d*. Also *connect*(*I*′, *I*) = *o*_1_ + *g*_2_ = *I*′.*b* − *I*.*a* + *I*.*c* − *I*′.*d* ≥ *I*.*c* + *I*.*b* − *I*.*a* − *I*′.*d* = *I*.*d* − *I*′.*d*.
3. **Case** *I*.*b* > *I*′.*b, I*.*a* ≤*I*′.*b, I*.*c* ≤*I*′.*d* (Fig. 5): Supposing wlog that *o*_1_ > *o*_2_, delete *S*_2_[*I*′.*d* + 1, *I*′.*d* + *o*_1_ − *o*_2_], and match *S*_1_[*I*′.*b* + 1, *I*.*b*] and *S*_2_[*I*′.*d* + *o*_1_ − *o*_2_ + 1, *I*.*d*]. This has edit cost *o*_1_ − *o*_2_ and *connect*(*I*′, *I*) = *o*_1_ − *o*_2_.
4. **Case** *I*.*b* > *I*′.*b, I*.*a* ≤*I*′.*b, I*.*c* > *I*′.*d* (Fig. 5): We delete *S*_2_[*I*′.*d* + 1, *I*′.*d* + *o*_1_ + *g*_2_] and match *S*_1_[*I*′.*b* + 1, *I*.*b*] with *S*_2_[*I*′.*d* + *o*_1_ + *g*_2_ + 1, *I*.*d*]. This has edit cost *o*_1_ + *g*_2_ and *connect*(*I*′, *I*) = *o*_1_ + *g*_2_.
5. **Case** *I*.*a* > *I*′.*b, I*.*c* > *I*′.*d* (Fig. 6): Supposing wlog *g*_2_ ≥*g*_1_, match with substitutions or unsupported exact matches *S*_1_[*I*′.*b* + 1, *I*′.*b* + *g*_1_] and *S*_2_[*I*′.*d* + 1, *I*′.*d* + *g*_1_]. Delete the substring *S*_2_[*I*′.*d* + *g*_1_ + 1, *I*.*c* − 1]. Finally, match *S*_1_[*I*.*a, I*.*b*] and *S*_2_[*I*.*c, I*.*d*]. The edits consist of *g*_1_ of substitutions or unsupported exact matches and *g*_2_ − *g*_1_ deletions, which is *g*_2_ edits in total. Also, *connect*(*I*′, *I*) = max*{g*_1_, *g*_2_*}* = *g*_2_.

**Fig. 4:**
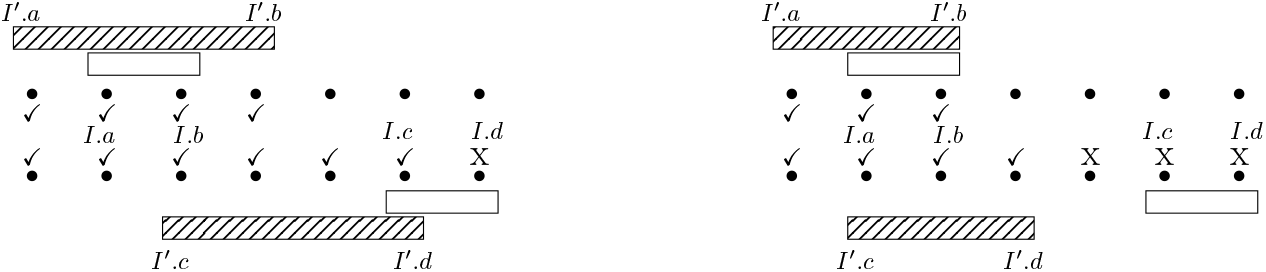
Cases in Proof of Lemma 1. The ✓symbol indicates symbols processed prior to considering *I*. (Left) Case *I*′, *b* ≥ *I*.*b* and *I*′.*d* ≥ *I*.*c* (Right) *I*′.*b* ≥ *I*.*b* and *I*′.*d* < *I*.*c*.

**Fig. 5:**
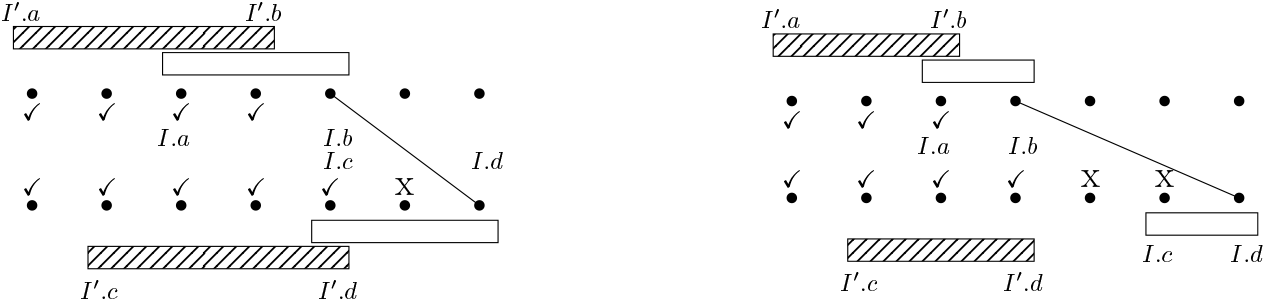
Cases in Proof of Lemma 1. (Left) Case *I*.*b* > *I*′.*b, I*.*a* ≤ *I*′.*b, I*.*c* ≤ *I*′.*d*. (Right) Case *I*.*b* > *I*′.*b, I*.*a* ≤ *I*′.*b, I*.*c* > *I*′.*d*.

**Fig. 6:**
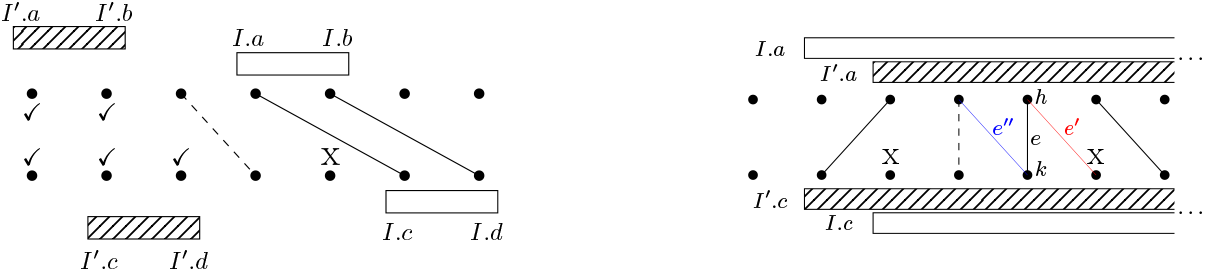
(Left) Case *I*.*a* > *I*′.*b, I*.*c* > *I*′.*d*. (Right) Anchors *I* and *I*′ are incomparable. The current alignment is shown with black solid and dashed edges. To remove *I*′ we sweep from right-to-left, replacing edges not supported by *I* with edges supported by *I*. Here, *e* = (*S*_1_[*h*], *S*_2_[*k*]) is not supported by *I* and will be replaced with *e*′ = (*S*_1_[*h*], *S*_2_[*I*.*c* + *h* − *I*.*a*]) (in red), which is supported by *I*.

Continuing this process until 𝒜_*right*_, all symbols in *S*_1_ and *S*_2_ become included in the alignment. □

We delay the details of Lemma 2‘s proof to Section 4.1.

#### Lemma 2.

*For a set of anchors 𝒜, optimal chaining cost under strict precedence* ≤ *anchored edit distance*.

*Proof*. We start with an arbitrary alignment ℳ supported by *A*. We will show in Lemma 3 how to obtain a subset ℬ ⊆ 𝒜 totally ordered under strict precedence and supporting an alignment ℳ′ where *EDIT* (ℳ′) ≤ *EDIT* (ℳ). We will then show in Lemma 4 that the edit cost of ℳ′ is greater or equal to the edit cost of the alignment ℳ_*G*_ given by the greedy algorithm on ℬ. Finally, in Lemma 5 we show that the co-linear chaining cost of ℬ is equal to the edit cost of ℳ_*G*_. Combining, we have *EDIT* (ℳ) ≥ *EDIT* (ℳ) ≥ *EDIT* (ℳ_*G*_) = the co-linear chaining cost on ℬ ≥ optimal co-linear chaining cost under strict precedence for 𝒜. The result follows from the fact that *EDIT* (ℳ) equals the anchored edit distance when ℳ is an optimal alignment for 𝒜. □

### 4.1 Details of Lemma 2 Proof

We apply Algorithm (i) followed by Algorithm (ii) to convert a supporting set of anchors 𝒜for ℳinto the totally ordered subset of anchors ℬsupporting ℳ′. Note that these algorithms are only for the purpose of the proof. Moving forward, we call an edge *e* = (*S*_1_[*h*], *S*_2_[*k*]) contained but not supported by *I* if *h* ∈ [*I*.*a, I*.*b*] or *k* ∈ [*I*.*c, I*.*d*] and *h* − *I*.*a* ≠ *k* − *I*.*c*. We define for *e* the two edges *e*′ = (*S*_1_[*h*], *S*_2_[*I*.*c* + *h*− *I*.*a*]) and *e*′′ = (*S*_1_[*I*.*a* + *k*− *I*.*c*], *S*_2_[*k*]), which are supported by *I*.

#### Algorithm (i). Algorithm for removing incomparable anchors

Let *I* and *I*′ be two incomparable anchors under weak precedence (Fig 6). The anchor that has the rightmost supported solid edge will be the anchor we keep. Suppose wlog it is *I*. Working from right-to-left, starting with that rightmost edge, for any edge *e* that is contained but not supported by *I*, we replace *e* with the rightmost of *e*′ and *e*′′. Note that at least one side of every edge supported by *I*′ is within an interval of *I*. Hence, all edges supported by *I*′ are eventually replaced. We then remove *I*′. This algorithm is repeated until a total ordering under weak precedence is possible.

#### Algorithm (ii). Algorithm for removing anchors with nested intervals

Consider two anchors *I* and *I*′ where wlog *I*′ has an interval nested in one of the intervals belonging to *I*. Let *e*_*R*_ be the rightmost edge supported by *I*. Working from right-to-left, we replace any edge *e* to the left of *e*_*R*_ that is contained but not supported by *I* with the rightmost of *e*′ and *e*′′. Next, working from left-to-right, we replace any edge *e* to the right of *e*_*R*_ that is contained but not supported by *I* with the leftmost of *e*′ and *e*′′. These procedures combined will replace all edges supported by *I*′ with those supported by *I*. We repeat this until there are no two nested intervals amongst all remaining anchors. Finally, remove all anchors that do not support any edge. We call such an anchor chain where every anchor supports at least one edge *minimal*.

##### Lemma 3.

*EDIT* (ℳ′) ≤ *EDIT* (*M*).

*Proof*. For Algorithm (i), suppose we are replacing an edge *e* not supported by the anchor *I*, the anchor we wish to keep. Suppose wlog that *e*′ is the rightmost of *e*′ and *e*′′, so we replace *e* with *e*′. Because the edge immediately to the right of *e* is also aligned with *I*, deleting *S*_2_[*k*] and matching *S*_2_[*I*.*c* + *h I*−.*a*], does not require modifying any additional edges. If *e* was a solid edge the edit cost is unaltered, since the total number of deletions and matches is unaltered. If *e* was a dashed edge, replacing *e* with *e*′ converts a substitution or unsupported exact match at *S*_2_[*k*] to a deletion, and removes a deletion at *S*_2_[*I*.*c* + *h − I*.*a*], decreasing the edit cost by 1. The same arguments hold for Algorithm (ii) when we replace edges from right-to-left. In Algorithm (ii) when we process edges from left-to-right, since any edges to left of the edge *e* being replaced are supported by *I*, replacing *e* with the leftmost of *e*′ and *e*′′ does not require modifying any additional edges. Again, if *e* is solid, the edit cost is unaltered, and if *e* is dashed, the edit cost is decreased by 1. □

##### Lemma 4.

*The greedy algorithm described in the proof of Lemma 1 produces an optimal alignment for a ‘minimal’ anchor chain under strict precedence*.

*Proof*. Proof is similar to proof of Lemma 3 and is deferred to Appendix 3.□

##### Lemma 5.

*For an anchor chain under strict precedence, the edit cost of the alignment produced by the greedy algorithm described in the proof of Lemma 1 is equal to the chaining cost*.

*Proof*. This follows from induction on the number of anchors processed, using the same arguments used in the proof of Lemma 1. However, only *I*′.*b* = *I*.*b* needs to be considered in Cases 1 and 2 leading to equality in these cases.□

## 5 Implementation

In multi-dimensional RmQs, *O*(*n*log^*d*−1^ *n*) storage requirement and irregular memory access during a query can limit their efficacy in practice [4]. We can take advantage of two observations to design a more practical algorithm. First, if sequences are highly similar, their edit distance will be relatively small. Hence the anchored edit distance, denoted in this section as *OPT*, will be relatively small for MUM or MEM anchors. Second, if the sequences are dissimilar, then the number of MUM or MEM anchors, *n*, will likely be small. These observations allow us to design an alternative algorithm (Algorithm 1) that requires *O*(*n*) worst-case space and *O*(*n·OPT* + *n* log *n*) average-case time over all possible inputs where *n* ≤max (|*S*_1_|, |*S*_2_|), i.e., the number of anchors is less than the longer sequence length (proof is deferred to Appendix 3). This property always holds when the anchors are MUMs and is typically true for MEMs as well. This makes the algorithm presented here a practical alternative.

As before, 𝒜 let be the initial (possibly unsorted) set of anchors, but with 𝒜 𝒜_*left*_ = 𝒜 [1] and 𝒜_*right*_ = 𝒜 [*n*]. We assume wlog |*S*_1_|≥|*S*_2_|. We begin by sorting anchor set 𝒜 by the component 𝒜[·].*a* and making a guess for the optimal solution, *B* (Algorithm 1). The value *B* is used at every step to bound the range of 𝒜[·].*a* values that need to be examined. This bounds the number of anchors that need to be considered (on average). If *C*[*n*] is greater than our current guess *B* after processing all *n* anchors, we update our guess to *B*_2_ · *B*.

Extending the above pseudo-code to enable semi-global chaining, i.e., free anchor gap on both ends of reference sequences, is also simple. In each *i*-loop, the connection to anchor 𝒜 _*left*_ must be always considered, and for last iteration when *i* = *n, j* must be set to 1. Second, a revised cost function must be used when connecting to either 𝒜_*left*_ or 𝒜_*right*_ where a gap penalty is used only for anchor gap over the query sequence. The experiments in the next section use an implementation of this algorithm.

## 6 Evaluation

There are multiple open-source libraries/tools that implement edit distance computation. Edlib (v1.2.7) [28] uses Myers’s bit-vector algorithm [20] and Ukkonen’s banded algorithm [29], and is known to be the fastest implementation currently.

### Algorithm 1: *O*(*OPT · n* + *n* log *n*) average-case algorithm.

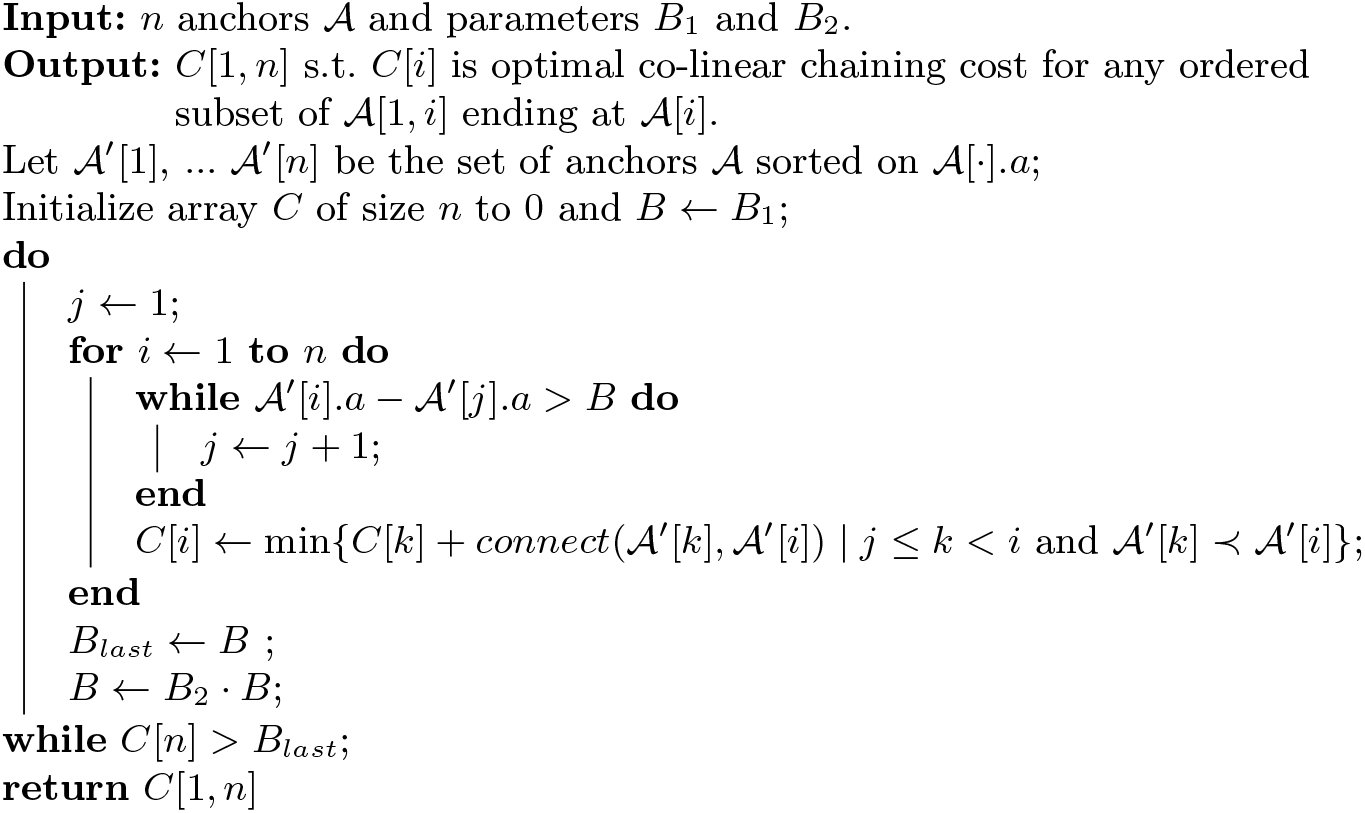

In this section, we aim to show that: (i) the proposed algorithm as well as existing chaining methods achieve significant speedup compared to computing exact edit distance using Edlib, and (ii) in contrast to existing chaining methods, our implementation consistently achieves high Pearson correlation (> 0.90) with edit distance while requiring modest time and memory resources.

We implemented Algorithm 1 in `C++`, and refer to it as ChainX. The code is available at https://github.com/at-cg/ChainX. Inputs are a target string, query strings, comparison mode (global or semi-global), anchor type preferred, i.e., maximal unique matches (MUMs) or maximal exact matches (MEMs), and a minimum match length. We include a pre-processing step to index target string using the same suffix array-based algorithm [31] used in Nucmer4 [18]. Chaining costs computed using ChainX for each query-target pair are provably-optimal.

### Existing co-linear chaining implementations

Co-linear chaining has been implemented previously as a stand-alone utility [2,22] and also used as a heuristic inside widely used sequence aligners [5,14,18]. Out of these, Clasp (v1.1), Nucmer4 (v4.0.0rc1) and Minimap2 (v2.22-r1101) tools are available as opensource, and used here for comparison purpose. Unlike our algorithm where the optimization problem involves minimizing a cost function, these tools execute their respective chaining algorithms using a score maximization objective function. Clasp, being a stand-alone chaining method returns chaining scores in its output, whereas we modified Minimap2 and Nucmer4 to print the maximum chaining score for each query-target string pair, and skip subsequent steps. To enable a fair comparison, all methods were run with single thread and same minimum anchor size 20. Accordingly, ChainX, Clasp and Nucmer4 were run with MUMs of length ≥20, and Minimap2 was configured to use minimizer *k*-mers of length 20. For these tests, we made use of an Intel Xeon Processor E5-2698 v3 processor with 32 cores and 128 GB RAM. All tools were required to match only the forward strand of each query string. ChainX and Clasp are both exact solvers of co-linear chaining problem, but use different gap-cost functions. Clasp only permits non-overlapping anchors in a chain, and supports two cost functions which were referred to as *sum-of-pair* and *linear* gap cost functions in their paper [22]. We tested Clasp with both of its gap-cost functions, and refer to these two versions as Clasp-sop and Clasp-linear respectively. Both the versions solve co-linear chaining using RmQ data structures, requiring *O*(*n* log^2^ *n*) and *O*(*n* log *n*) time respectively. Both require a set of anchors as input, therefore, we supplied the same set of anchors, i.e., MUMs of length ≥ 20 as used by ChainX. Minimap2 and Nucmer4 use co-linear chaining as part of their seed-chain-extend pipelines. Both Minimap2 and Nucmer2 support anchor overlaps in a chain, as well as penalize gaps using custom functions. However, both these tools employ heuristics (e.g., enforce a maximum gap between adjacent chained anchors) for faster processing which can result in suboptimal chaining scores.

### Runtime and memory comparison

We downloaded the same set of query and target strings that were used for benchmarking in Edlib paper [28]^2^. These test strings, ranging from 10 kbp to 5000 kbp in length, allowed us to compare tools for end-to-end global sequence comparisons as well as semi-global comparisons at various degrees of similarity levels. To test end-to-end comparisons, the target string had been artificially mutated at various rates using mutatrix (https://github.com/ekg/mutatrix), whereas for the semi-global comparisons, a substring of the target string had been sampled and mutated. Table 1 presents runtime and memory comparison of all tools. Columns of the table are organized to show tools in three categories: edit distance solver (Edlib); optimal co-linear chaining solvers (ChainX, Clasp-sop, Clasp-linear); and heuristic implementations (Nucmer4, Minimap2). We make the following observations here. First, chaining methods (both optimal and heuristic-based) are significantly faster than Edlib in most cases, and we see up to three orders of magnitude speedup. Second, within optimal chaining methods, Clasp-sop’s time and memory consumption increases quickly with increase in count of anchors, which is likely due to irregular memory access and storage overhead of its algorithm that uses a 2d-RmQ data structure. Finally, we note that Minimap2 and Nucmer4 are often faster than exact algorithms during global string comparisons due to their fast heuristics.

**Table 1:**
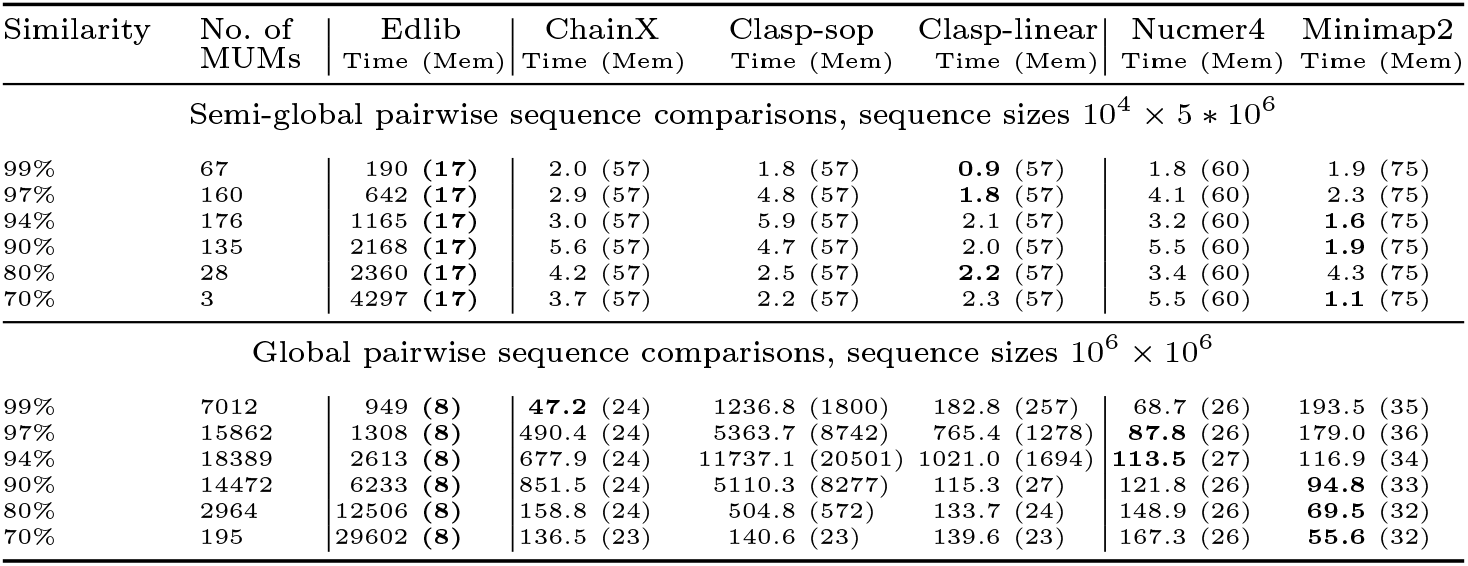
Runtime and memory usage comparison of edit distance solver Edlib and co-linear chaining methods ChainX, Clasp, Nucmer4 and Minimap2. Runtime is measured in milliseconds across the columns, and memory usage (Mem) is noted in MBs. In this experiment, ChainX, Clasp-sop, Clasp-linear and Nucmer4 used maximal unique matches (MUMs) of length ≥ 20 as input anchors, while Minimap2 used fixed-length minimizer *k*-mers of size 20.

All tools (except Edlib) use an indexing step such as building a *k*-mer hash table (Minimap2) or computing suffix array (ChainX, Clasp-sop, Clasp-linear, Nucmer4). Indexing time was excluded from reported results, and was found to be comparable. For instance, in the case of semi-global comparisons, ChainX, Nucmer4, Minimap2 required 590 ms, 736 ms, 236 ms for index computation respectively.

### Correlation with edit distance

We checked how well the chaining cost (or score) correlates with edit distance. We use absolute value of Pearson correlation coefficients for a comparison. In this experiment, we simulated 100 query strings within three similarity ranges: 90 − 100%, 80 − 90% and 75 − 80%. Table 2 shows the correlation achieved by all the tools. Here we observe that ChainX and Clasp-sop are more consistent in terms of maintaining high correlation across all similarity ranges. Between the two, ChainX was shown to offer superior scalability in terms of runtime and memory usage (Table 1). Hence, ChainX can be useful in practice when good performance and accuracy is desired across a wide similarity range.

**Table 2:**
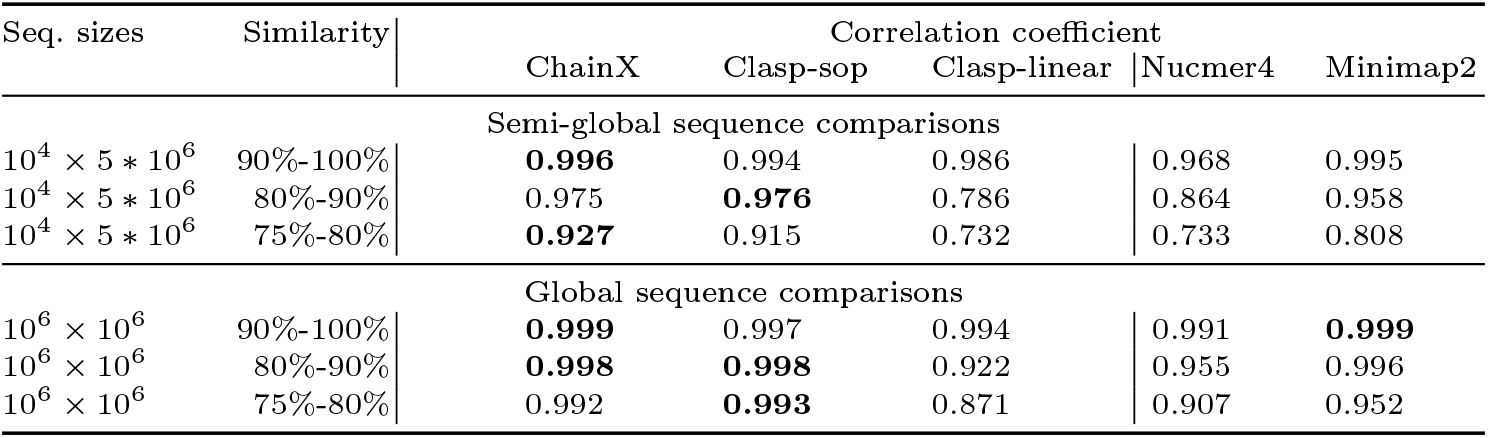
Absolute Pearson correlation coefficients of chaining costs (or scores) computed by various methods with the corresponding edit distances. 100 query strings were simulated and matched to the target string within each similarity range.

| Seq. sizes | Similarity | Correlation coefficient |  |  |  |  |
| --- | --- | --- | --- | --- | --- | --- |
|  |  | ChainX | Clasp-sop | Clasp-linear | Nucmer4 | Minimap2 |
| Semi-global sequence comparisons |  |  |  |  |  |  |
| $10^4 \times 5 * 10^6$ | 90%-100% | **0.996** | 0.994 | 0.986 | 0.968 | 0.995 |
| $10^4 \times 5 * 10^6$ | 80%-90% | 0.975 | **0.976** | 0.786 | 0.864 | 0.958 |
| $10^4 \times 5 * 10^6$ | 75%-80% | **0.927** | 0.915 | 0.732 | 0.733 | 0.808 |
| Global sequence comparisons |  |  |  |  |  |  |
| $10^6 \times 10^6$ | 90%-100% | **0.999** | 0.997 | 0.994 | 0.991 | **0.999** |
| $10^6 \times 10^6$ | 80%-90% | **0.998** | **0.998** | 0.922 | 0.955 | 0.996 |
| $10^6 \times 10^6$ | 75%-80% | 0.992 | **0.993** | 0.871 | 0.907 | 0.952 |

### Effect of anchor type and minimum match length

How many anchors are given as input will naturally affect the performance and output quality of a chaining algorithm. We tested runtime and correlation with edit distance achieved by ChainX while varying the anchor type (MUMs/MEMs) and minimum match-length *l*_*min*_ parameter (Table 3). When MUMs are used as anchors, we observe good scalability, and lowering *l*_*min*_ from 20 to 10 improves the correlation, but the correlation saturates afterwards. This is because very short exact matches will unlikely be unique and won’t be selected as MUMs. However, when MEMs are used as anchors, correlation continues to improve with decreasing minimum length parameter, however, runtime grows exponentially. Excessive count of anchors improves the correlation but then anchor chaining becomes computationally demanding.

**Table 3:**
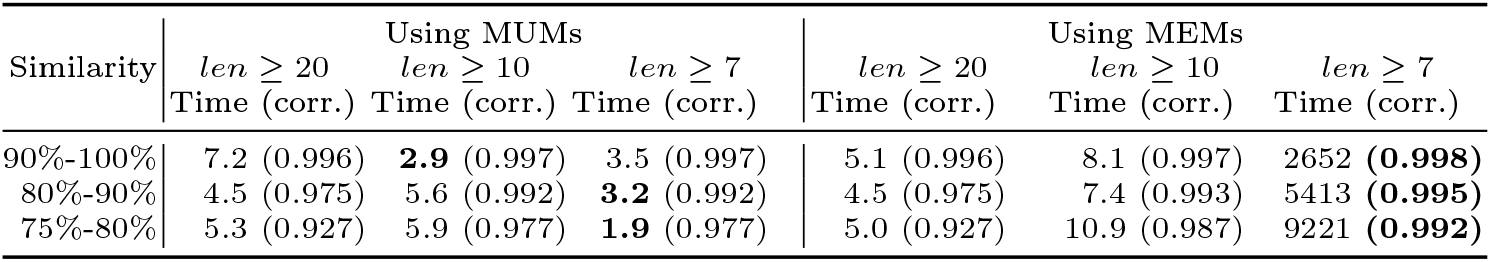
Effect of anchor pre-computation method on the performance of ChainX. Total runtime to do 100 pairwise semi-global sequence comparisons (sequence size: 10^4^*×* 5 *10^6^) is measured in seconds, and correlation (corr.) with the corresponding edit distances is computed using Pearson correlation coefficient.

## 7 Conclusions

This work provides new algorithms for co-linear chaining, a fundamental problem in bioinformatics. Variants of this technique have been regularly used in alignment tools since four decades [32]. We addressed an open problem pertaining to the general case of this problem which allows anchor overlaps and penalizes gap cost between adjacent chained anchors. The proposed algorithms for multiple versions of this problem are provably-optimal and efficient, and can be incorporated in read mappers. We also discussed a new cost function for the co-linear chaining problem that enabled us to establish the first mathematical link between co-linear chaining and the edit distance problem. This result is a useful addition to a prior result [16] where a connection between the co-linear chaining problem and the longest common subsequence problem was made.

## Acknowledgements

This research is supported in part by the U.S. National Science Foundation (NSF) grants CCF-1704552, CCF-1816027, CCF-2112643, and funding from the Indian Institute of Science.

## Appendix

### 1 Algorithm Pseudocode

#### Algorithm 2: For co-linear chaining with overlaps and gap costs

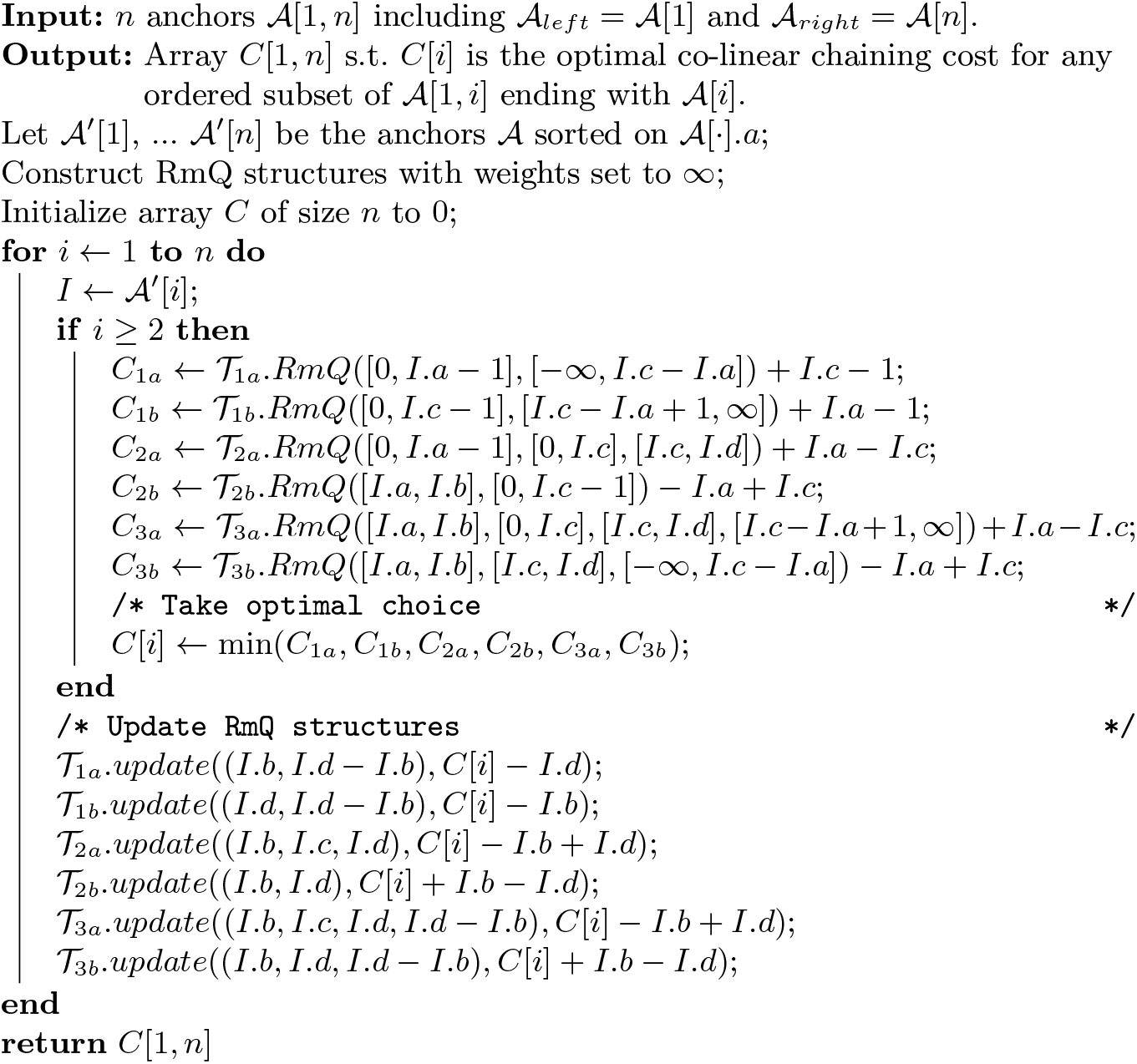

### 2 Modifications for weak precedence and fixed-length anchors

We first consider the case of weak precedence. In Case 3.a the anchor end *I*′.*d* can be positioned arbitrarily to the right of *I*.*c*. Moreover, since by the first dimension of the RmQ there is positive overlap in *S*_1_ and by the fourth dimension there is greater overlap in *S*_2_, we know that *I*′.*d* ≥ *I*.*c*. Hence, we can then remove the third dimension from the RmQ. The query will then be of the form (*I*′.*b, I*′.*c, I*′.*d*− *I*′.*b*) ∈ [*I*.*a*, ∞ ] × [0, *I*.*c*] *×* [*I*.*c*− *I*.*a* + 1, ∞ ]. In Case 3.b, where there is greater or equal overlap in *S*_1_, we can similarly ignore *I*′.*b*, but in order to match our definition of weak precedence we must also ensure *I*′.*c* ∈ [0, *I*.*c* −1] (this is unnecessary for *I*′.*a* in Case 3.a as the strictly greater overlap in *S*_2_ ensures *I*′.*a* < *I*.*a*). We modify the query to be of the form (*I*′.*c, I*′.*d, I*′.*d* − *I*′.*b*) ∈ [0, *I*.*c*− 1] *×* [*I*.*c*, ∞] *×* [−∞, *I*.*c − I*.*a*]. Since each RmQ has at most three dimensions, the total time complexity can be brought down to *O*(*n* log^3^ *n*).

In the case of fixed-length anchors, the RmQ for Case 2.a. can be made (*I*′.*b, I*′.*d*) ∈ [0, *I*.*a* −1] *×* [*I*.*c, I*.*d*]. The modifications for Cases 3.a and 3.b are more involved. We keep a pointer *p*_*a*_ to indicate the current *a* value of the interval, initially setting *p*_*a*_ = 𝒜 [1].*a*. Conceptually, before processing anchor *I* we increment *p*_*a*_ from its previous position to *I*.*a*. If for some anchor *I*′ the end *I*′.*b* is passed by *p*_*a*_, we update the points associated with *I*′ in 𝒯_3*a*_ and 𝒯_3*b*_ to have the the weight. This eliminates the need to use a range query to check *I*′.*b* ∈ [*I*.*a, I*.*b*], since any points not within that range are effectively removed from consideration. Hence, we can reduce the query for Case 3.a. (overlap in *S*_2_ greater than overlap in *S*_1_) to (*I*′.*c, I*′.*d* − *I*′.*b*) ∈ [0, *I*.*c*] *×* [*I*.*c − I*.*a* + 1, ∞]. and the query for Case 3.b (overlap in *S*_1_ greater or equal to overlap in *S*_2_) to (*I*′.*d, I*′.*d* − *I*′.*b*) ∈ [*I*.*c, I*.*d*] *×* [−∞, *I*.*c* − *I*.*a*]. To avoid the |*S*_1_| time complexity, the anchors that would be encountered while incrementing *p*_*a*_ can be found by looking at which anchors have *b* values between the previous *p*_*a*_ value and *I*.*a*. Because each update requires *O*(log^2^ *n*) time, these updates cost time *O*(*n* log^2^ *n*) in total.

### 3 Missing proofs

#### Lemma 4.

*For a minimal anchor chain under strict precedence, the greedy algorithm described in Lemma 1 produces an optimal alignment*.

*Proof*. This follows from an exchange argument. Suppose there exists an optimal alignment ℳ on the anchor chain ℬ that is not the same as the alignment ℳ_*G*_ produced by the greedy algorithm. As we process the edges from left-to-right, consider when the first discrepancy in the edges is found, the leftmost edges *e* = (*S*_1_[*h*], *S*_2_[*k*]) in ℳand *e*_*G*_ = (*S*_1_[*h*_*G*_], *S*_2_[*k*_*G*_]) in ℳ_*G*_ that are not equal. Let *e*_*prev*_ be the previous edge on which ℳ the and ℳ_*G*_ coincided. We claim *e* can be replaced with *e*_*G*_ without increasing the edit cost.

– Case: *e*_*G*_ is solid and left of *e*. Then *e* can clearly be exchanged for *e*_*G*_ with no increase in edit cost.
– Case: *e*_*G*_ is solid and not left of *e*. We assume WLOG *k* < *k*_*G*_ (Figure 7). Suppose *e*_*G*_ is supported by an anchor *I*. We claim that (i) *h ≥ h*_*G*_, and (ii) there exists an edge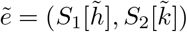 in ℳ to the right of *e* and supported by *I*. Note that 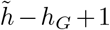 solid edges can be obtained by matching 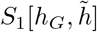] to 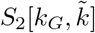, and that this alignment has at most the edit cost incurred within the intervals 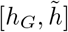 in *S*_1_ and 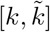 in *S*_2_ by *ℳ*. Thanks to (i) and (ii), swapping all of the edges in ℳ inside these intervals with the 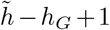 edges supported by *I* mentioned above can be done without effecting edges outside those intervals. The proofs for claims (i) and (ii) are shown below.
  i. *h* ≥*h*_*G*_: Suppose to the contrary, that *h* < *h*_*G*_. Let *e*_*prev*_ = (*S*_1_[*h*_*prev*_], *S*_2_[*k*_*prev*_]). Since *h*_*prev*_ < *h* < *h*_*G*_ and *k*_*prev*_ < *k* < *k*_*G*_, this would imply that there are deletions (x’s) at both *S*_1_[*h*] and *S*_2_[*k*] in ℳ_*G*_. However, the greedy algorithm would instead create an edge including *S*_1_[*h*] or *S*_2_[*k*] (or both), causing there to be an edge to the left of *e*_*G*_ and to the right of *e*_*prev*_. This contradicts our assumption that *e*_*G*_ is the first edge in ℳ_*G*_ to the right of *e*_*prev*_.
  ii. *An edge* 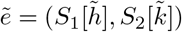 *exists in ℳ to the right of e and supported by I:* We claim *e*_*G*_ is the leftmost edge supported by *I* in ℳ_*G*_. Otherwise, since only consecutive edges supported by an anchor are taken in the greedy algorithm (Cases 3, 4, 5 in proof of Lemma 1), *e*_*prev*_ = (*S*_1_[*h*_*G*_ − 1], *S*_2_[*k*_*G*_ − 1]). Then *e*_*prev*_ was also in ℳ by our assumption that ℳ_*G*_ and ℳ agreed until *e*_*G*_. However, *e*_*prev*_ = (*S*_1_[*h*_*G*_ − 1], *S*_2_[*k*_*G*_ − 1]) would cross or share a vertex with *e* = (*S*_1_[*h*], *S*_2_[*k*]) in ℳ since *h*_*G*_ − 1 < *h* and *k* ≤ *k*_*G*_ − 1, a contradiction. Now, if no other edges supported by *I* are in ℳ, then ℬ is not minimal. Hence, we can assume that an edge 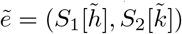 exists in ℳ to the right of *e* and supported by *I*.
– Case: *e*_*G*_ is a dashed edge. Since dashed edges are only created in Case 5 of the proof of Lemma 1 (Figure 6), *e*_*prev*_ = (*S*_1_[*h*_*G*_ −1], *S*_2_[*k*_*G*_ − 1]) must have been either the rightmost supported edge for some anchor, or another dashed edge. Additionally, no other supported edges are possible that include either *S*_1_[*h*] or *S*_2_[*k*]. Here the dashed edge *e*_*G*_ is optimal and swapping *e* with *e*_*G*_ will reduce the edit cost.□

**Fig. 7:**
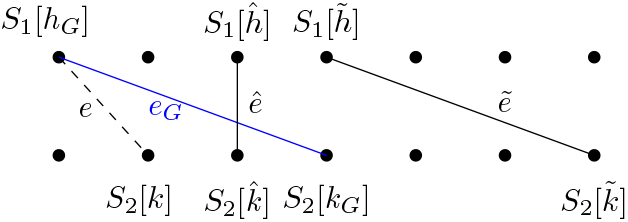
Consider a greedy alignment ℳ_*G*_ and optimal alignment ℳ where the leftmost difference is *e* and *e*_*G*_ (in blue). Here, *ê* prevents swapping *e* and *e*_*G*_ without modifying additional edges in ℳ.

#### Lemma 6.

*Algorithm 1 runs in O*(*n · OPT* + *n* log *n*) *average-case time over all inputs where n* ≤ max(|*S*_1_|, |*S*_2_|).

*Proof*. The *n* log *n* term is from the sorting the anchors. To analyze the second portion of the algorithm, we first let *X*_*h,j*_ be 1 if 𝒜 [*h*].*a* is located at index *j* in *S*_1_. Under the assumption of a random placement of anchors, 𝔼 [*X*_*h,j*_] = 1*/*|*S*_1_|. Let *X*_*i*_ be the number of anchors, 𝒜 [*h*], where 𝒜 [*h*].*a* ∈ [𝒜 [*i*].*a* − *B, 𝒜* [*i*].*a* − 1]. We have that 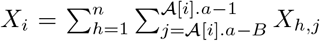. Letting *X* be the total number of anchors processed, 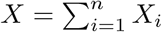 and

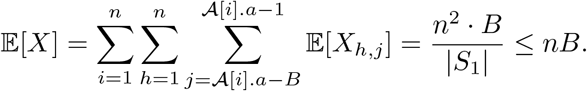

The total expected time is a constant factor from 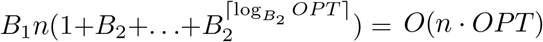.□

## Footnotes

1 *Õ*(*·*) hides poly-logarithmic factors.

2 https://github.com/Martinsos/edlib/tree/master/test_data

